# Visual working memory is independent of the cortical spacing between memoranda

**DOI:** 10.1101/216341

**Authors:** William J. Harrison, Paul M. Bays

## Abstract

The sensory recruitment hypothesis states that visual short term memory is maintained in the same visual cortical areas that initially encode a stimulus’ features. Although it is well established that the distance between features in visual cortex determines their visibility, a limitation known as crowding, it is unknown whether short term memory is similarly constrained by the cortical spacing of memory items. Here we investigated whether the cortical spacing between sequentially presented memoranda affects the fidelity of memory in humans (of both sexes). In a first experiment, we varied cortical spacing by taking advantage of the log-scaling of visual cortex with eccentricity, sequentially presenting memoranda in peripheral vision along either the radial or tangential visual axis with respect to the fovea. In a second experiment, we sequentially presented memoranda either within or beyond the critical spacing of visual crowding, a distance within which visual features cannot be perceptually distinguished due to their nearby cortical representations. In both experiments and across multiple measures, we found strong evidence that the ability to maintain visual features in memory is unaffected by cortical spacing. These results indicate that the neural architecture underpinning working memory has properties inconsistent with the known behaviour of sensory neurons in visual cortex. Instead, the dissociation between perceptual and memory representations supports a role of higher cortical areas, such as posterior parietal or prefrontal regions, or may involve an as yet unspecified mechanism in visual cortex in which stimulus features are bound to their temporal order.

**Significance Statement:** Although much is known about the resolution with which we can remember visual objects, the cortical representation of items held in short term memory remains contentious. A popular hypothesis suggests that memory of visual features is maintained via the recruitment of the same neural architecture in sensory cortex that encodes stimuli. We investigated this claim by manipulating the spacing in visual cortex between sequentially presented memoranda such that some items shared cortical representations more than others, while preventing perceptual interference between stimuli. We found clear evidence that short term memory is independent of the intra-cortical spacing of memoranda, revealing a dissociation between perceptual and memory representations. Our data indicate that working memory relies on different neural mechanisms from sensory perception.

## Introduction

Although a focus of research for decades, the neural basis of working memory storage is still disputed (Serences, 2016; Xu, 2017). Recent neuro-imaging studies have demonstrated that items in memory can be decoded from activity in human primary visual cortex (V1). Whereas the amplitude of the blood-oxygenation-level-dependent (BOLD) signal within V1 is not predictive of a remembered stimulus, patterns of activity across voxels can reliably predict memoranda (Harrison & Tong, 2009; Serences, Ester, Vogel, & Awh, 2009). For example, in a study by Harrison and Tong, observers viewed two sequentially presented oriented gratings and were cued to hold one item in memory so that they could later compare it with a test grating. These authors found that the remembered stimulus orientation could be decoded from patterns of activity within V1 during the retention interval. They concluded that visual cortex retains information about features in working memory. Similar studies have found that activity patterns within early visual cortex are specific to only the task-relevant feature of multi-feature objects (Serences et al., 2009) and that the precision of decoding diminishes with increasing numbers of memoranda (Emrich, Riggall, LaRocque, & Postle, 2013; Sprague, Ester, & Serences, 2014).

These findings among others have led some researchers to conclude that memory storage mechanisms are located within the sensory neural systems involved in processing the stimulus attributes, a proposal termed the *sensory recruitment hypothesis* (Emrich et al., 2013; Pasternak & Greenlee, 2005; Serences, 2016; Sreenivasan, Curtis, & D’Esposito, 2014). This hypothesis is appealing in part because visual cortex is thought to be one of the few brain areas with sufficient processing power to represent objects with the level of detail observed in short term memory (for a review, see Serences, 2016). However, it is not clear how visual cortex could maintain memory representations while simultaneously processing new incoming information, nor how the different perceptual experiences of seeing versus remembering are accounted for by this hypothesis.

In contradiction to the sensory recruitment hypothesis, Bettencourt and Xu (2016) found that target features could not be decoded from early visual cortex when distractors were presented during the memory retention period, but that such distractors had no impact on behavioural performance. Bettencourt and Xu could reliably decode activity within a region of parietal cortex to predict the target stimulus regardless of whether or not a distractor was presented, suggesting an important role of that area in short term memory. It remains contentious, therefore, whether visual cortex plays a necessary role in short term memory maintenance (Xu, 2017).

In the present study, we tested whether the fidelity with which memoranda are stored is affected by the neural resources available within early visual cortex, by varying the intra-cortical spacing of items. When items are presented simultaneously, in the absence of working memory demands, their intra-cortical spacing is the primary constraint on their perceptual discriminability. Nearby stimuli “crowd” each other, and the zone of crowding is determined by the distance between stimuli in retinotopic cortex (Pelli, 2008; Pelli & Tillman, 2008). Visual crowding occurs when the cortical spacing between visual objects prevents a distinct target representation in early visual cortex (Anderson, Dakin, Schwarzkopf, Rees, & Greenwood, 2012; J. Chen et al., 2014; Kwon, Bao, Millin, & Tjan, 2014; Pelli, 2008; van den Berg, Roerdink, & Cornelissen, 2010), or results in pooling of stimulus representations at later levels of the visual hierarchy (Freeman & Simoncelli, 2011). If short term memory of items presented in spatial isolation is maintained via the recruitment of the same sensory areas involved in the encoding of those features, then we should see worse memory performance for items that are closer together in visual cortex, and therefore share more neural resources, than for items with greater intra-cortical spacing.

## Materials and Methods

### Experiment 1 Overview

We investigated whether log-scaling of visual cortex affects short term memory by having observers remember three items on each trial arranged according to one of two spatial configurations, and, using a method of adjustment, report the orientation of the item indicated by a probe. Within a trial, items were aligned along either the tangential axis or the radial axis, and thus had greater or lesser intra-cortical spacing, respectively (Fig. 1A and 1B). Importantly, in each configuration, one item appeared at 10° eccentricity directly to the right of fixation, and so targets at this location were matched in all regards except for the intra-cortical spacing between memoranda within the same trial. We thus focus analyses only on target items at this location, although all locations were probed equally often so as to encourage participants to store all items in short term memory. Finally, we ensured our data were not confounded by perceptual interference (e.g. Yeshurun, Rashal, & Tkacz-Domb, 2015) by presenting items sequentially and with sufficient durations and interstimulus intervals to negate such perceptual effects.

**Figure 1.**
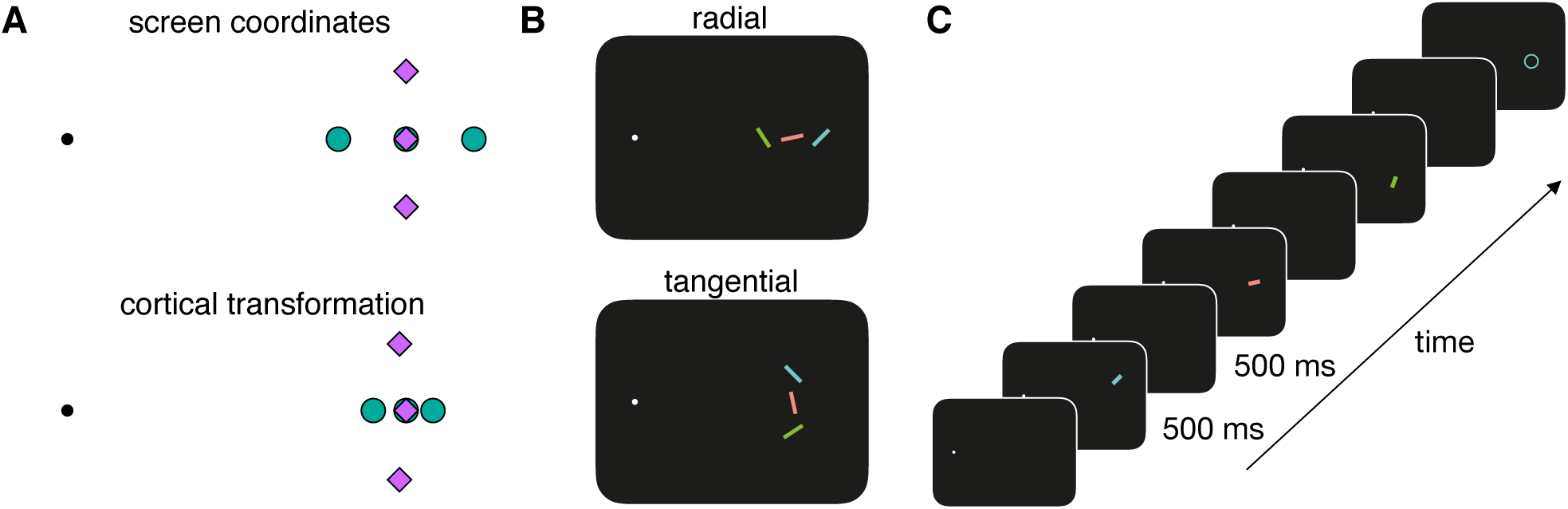
Experiment 1 design. A) Differences in cortical spacing in peripheral vision. The top row depicts the screen coordinates of stimuli in peripheral vision with respect to the point of fixation (black spot). The inter-item spacing following cortical transformation is shown in the bottom row. Such a cortical representation of space occurs in V1, which is hypothesised to maintain memory representations. Cortically transformed coordinates are normalised to the central target position. Green spots and purple diamonds represent radial and tangential spatial arrangements of stimuli, respectively. Note that, although stimuli are equally spaced in screen coordinates across conditions, radially arranged stimuli have less intra-cortical spacing than tangentially arranged stimuli. B) Stimulus design. Memoranda were randomly oriented coloured bars, presented sequentially along either the radial or tangential axis. Note that the centre stimulus in each condition occupies the same screen (and therefore cortical) location. C) Example trial sequence. Observers fixated a white spot while memoranda were presented in sequence. Following a delay after the presentation of the third item, a probe was shown matching the colour and location of one item chosen at random, cueing observers to move the mouse to report the remembered orientation of that item. A response bar appeared within the circle after the first mouse movement was detected, allowing observers to make their response using a method of adjustment.

#### Participants

10 people participated in Experiment 1 (mean age 24 ± 3.07; 5 male, 5 female). All had typical colour vision and normal or corrected-to-normal acuity and were naïve to the purposes of the experiment. All observers gave written informed consent and were paid £10 per hour for their participation. The study was approved by the University of Cambridge Psychology Research Ethics Committee.

#### Experimental Setup

Participants sat in a head and chin rest positioned 57 cm from an ASUS LCD monitor. The resolution of the monitor was 1920 × 1200 within an area that was 44.8 cm × 28 cm with no pixel interpolation. Stimulus colours were selected after measuring the luminance of each colour channel of the monitor with a spectrophotometer. Fixation was monitored online with an EyeLink 1000 (SR Research) recording at 500Hz, calibrated once before each testing session and re-calibrated as required throughout the experiment (see below). The experiment was programmed with the Psychophysics Toolbox Version 3 (Brainard, 1997; Pelli, 1997) and Eyelink Toolbox (Cornelissen, Peters, & Palmer, 2002) in MATLAB (MathWorks).

#### Stimuli

On each trial, three randomly oriented bars (2° × 0.2° of visual angle) were presented sequentially, and each was uniquely coloured red, green, or blue. Colours were matched in luminance (26.2 cd/m^2^) and the order in which they appeared as well as their screen position were randomised across trials. A white fixation spot was displayed in the centre of the screen throughout stimulus presentation and the memory delay period. All stimuli were presented on a black background (luminance < 1 cd/m^2^).

Within a trial, stimulus positions were arranged either tangentially or radially with respect to the point of fixation (Fig. 1B). In both conditions, one item was centred on the horizontal meridian, 10° right of fixation. In the radial condition, the two other items were positioned 2° left or right of the central item, such that they were arranged along the horizontal meridian. In the tangential condition, one item was positioned 2° above the central item, and the other was positioned 2° below the central item, such that they were arranged orthogonal to the horizontal meridian. Although never presented simultaneously, the inter-stimulus spacing meant that their positions did not overlap. The order in which a stimulus was presented at each position was randomised across trials.

#### Procedure

A typical trial sequence is shown in Figure 1C. At the start of each trial, an observer had to maintain fixation within a 2° region of the fixation spot for 500 ms for the trial to proceed. If fixation remained outside this region for more than 2 seconds, the eye tracker was re-calibrated. Once correct fixation was registered, there was an additional variable delay between 250 – 750 ms (uniformly distributed). Stimuli were then presented sequentially in either a tangential or radial arrangement (Fig. 1B). The stimulus duration and inter-stimulus interval were 500 ms. Following the offset of the third stimulus, there was a 500 ms delay period, after which a probe circle (2° diameter) appeared centred on the location previously occupied by one of the three items, cueing the observer to report the orientation of that item using the mouse. Once any movement of the mouse was recorded, a response bar replaced the probe circle and followed the orientation designated by the mouse position relative to the bar centre. The response bar had the same dimensions as the target item, but its orientation was randomised at the start of each response period and remained on screen until the observer clicked the mouse button to confirm their response.

During pilot testing with white stimuli, we noted that it was difficult to attribute clearly the probe circle to one memory item based on location alone, particularly for the radial condition. This is most likely due to the well-known compression of perceptual space in peripheral vision (McGraw & Whitaker, 1999; White, Levi, & Aitsebaomo, 1992), and so during Experiment 1 the probe circle and response bar also matched the colour of the target item.

Participants were informed that all items were equally likely to be the target. The target appeared equally as often across temporal order and location. There were 324 trials, consisting of 18 repetitions for each target combination (3 target locations for each of 2 spatial arrangements and 3 temporal orders).

If gaze position deviated by more than 2° from the fixation spot during stimulus presentation, the inter-stimulus interval, or the delay period, the message, “Don’t look away from the fixation point until it’s time to respond,” appeared for two seconds, and the trial restarted with newly randomised stimulus orientations. Each testing session took approximately one hour. After 50% of trials were completed, the observers were requested to take a short break, but were also instructed they could rest at other times as they required.

#### Experimental Design and Analyses

All comparisons in this experiment were within-subjects. Only trials in which the target item was positioned 10° to the right of fixation were analysed. For items at this location, we compared memory performance across radial and tangential conditions with two measures, collapsed across temporal order. We first analysed the variability of report errors by calculating the circular standard deviation of reports for each condition for each observer. These values were compared across conditions with a Bayesian t-test using JASP software (JASP Team, 2017). We used the default Cauchy prior width of 0.707, but all results reported below were robust to standard alternate prior widths. Alongside Bayes factors, we provide Student t-test results.

In a second analysis, we assessed whether there was an influence of intra-cortical spacing on observers’ reports using a probabilistic model of working memory performance. This was the “swap” model introduced by Bays et al. (2009), in which observers’ responses are attributed to a mix of noisy reports centred on the target orientation, noisy reports centred on non-target items, and a uniform lapse rate (see also Zhang & Luck, 2008). The details of this model have been described extensively elsewhere (for examples, see Bays et al., 2009; Gorgoraptis, Catalao, Bays, & Husain, 2011). The model has three free parameters: precision of reports, proportion of swap errors, and proportion of guesses. Parameters were estimated by maximum likelihood using code available online (http://www.paulbays.com/code/JV10; Bays et al., 2009).

We compared two versions of the model: a full model in which a separate set of parameters was used for radial and tangential conditions, and a restricted model in which a single set of parameters was used for both conditions. To compare which of the two models best described the data, we used the Akaike Information Criterion (AIC) summed across participants. To further test whether the models differentially accounted for the data, we submitted the differences in individuals’ AIC scores to Bayesian and Student t-tests.

### Experiment2 Overview

Experiment 2 was designed to ensure the physical spacing between memoranda would result in competing representations within primary visual cortex. For each participant in Experiment 2, we first measured the critical spacing of crowding, which is the area within which crowding occurs (Pelli & Tillman, 2008). We then tested observers’ memory for memoranda presented sequentially within versus beyond their critical spacing. Moreover, we tested whether there is a correlation between critical spacing and memory performance, which could arise if working memory is related to individual differences in cortical surface area (e.g. Schwarzkopf, Song, & Rees, 2011). To increase statistical power and to assess the correlation between critical spacing and memory performance, we greatly increased the sample size compared with Experiment 1.

Each participant first completed a crowding task in which we found the inter-item distance at which their ability to recognise a target reached threshold level, which we take as the critical spacing of crowding. A participant’s basic task was to identify the orientation of a bar surrounded by a circle, flanked on either side by distractors (Fig. 3A). Target and distractors were briefly presented in the upper peripheral visual field, and trial-by-trial variations in interitem spacing were controlled by an adaptive procedure. The participant reported the target orientation by clicking on one of three response options shown around the point of fixation (a three alternative forced-choice task). After finding their critical spacing, the participant then completed a memory experiment in which three randomly oriented bars were presented in sequence in one of two spatial configurations (Fig. 4A). Within each trial, memoranda were presented across a spatial range equal to either 0.75 or 1.5 times their critical spacing, corresponding to “crowded” and “uncrowded” conditions, respectively. As in Experiment 1, there was a common screen position for one item in each condition, and we analysed only memory performance for this stimulus position. Therefore, any differences in performance across conditions could only be driven by differences in intra-cortical spacing of memoranda.

**Figure 3.**
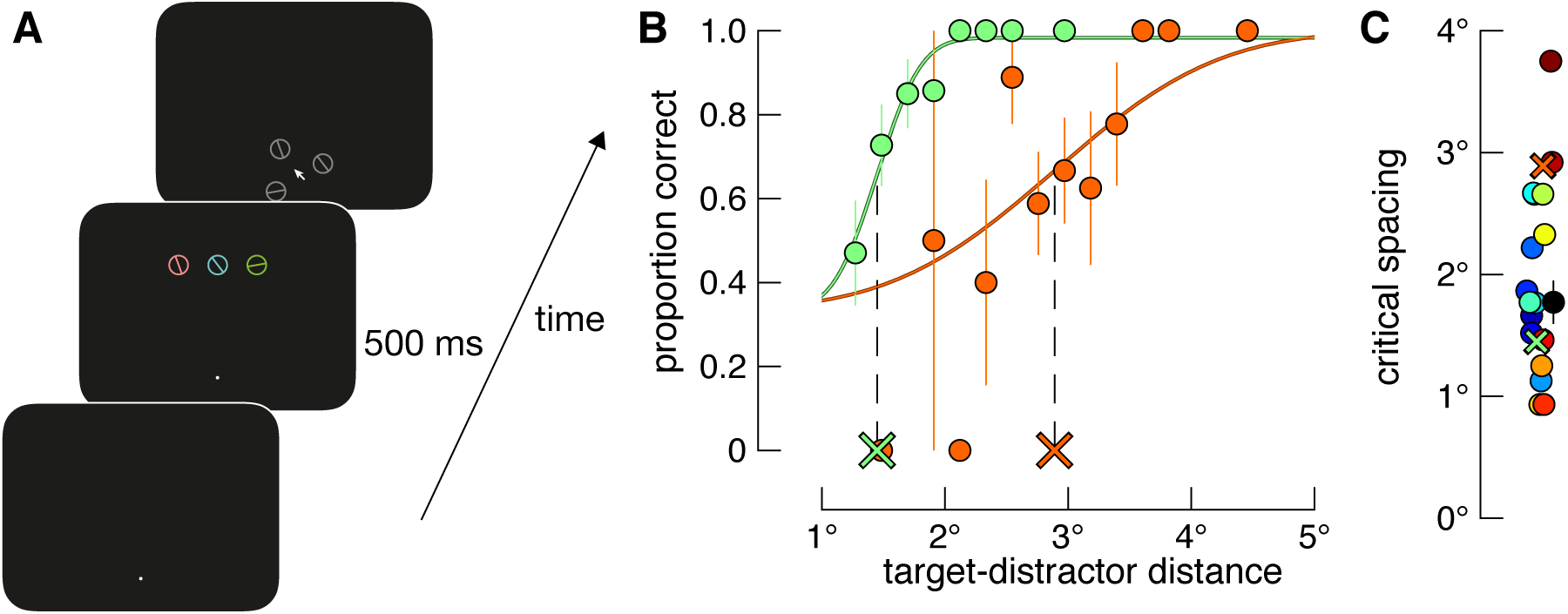
Design and results of the crowding task. A) Example trial sequence. After fixating a white spot, three stimuli were presented in the upper visual field. An observer’s task was to identify the orientation of the centre stimulus, and report its orientation by clicking on the matching stimulus in grey in the subsequent display. The distance between target and distractors on each trial was controlled via an adaptive procedure. B) Example results and psychometric functions. Differently coloured data show results for two differently performing observers. Solid lines show Weibull functions fit to each dataset. Dashed black lines and coloured X symbols show the midpoint of the function and corresponding critical spacing estimates, respectively, for each participant. C) Estimated critical spacing for 19 observers. The median critical spacing is shown as the black datum, while individual participants’ values are shown in various colours. Estimates corresponding to the psychometric functions in (B) are shown as X symbols. Data have been randomly jittered on the x-axis to minimise overlap. Error bars in all panels show ±1 SE.

Results from the memory experiment are shown in Figure 4B-E. We first compared observers’ report variability for the crowded and uncrowded conditions (Fig. 4B). These data are summarised as difference scores in Figure 4C. Rather than finding an effect of crowding on response standard deviation, a Bayesian paired-samples t-test found moderate evidence in favour of there being no difference between conditions (BF_01_ = 4.21; t(18) = 0.051, p = 0.96).

**Figure 4.**
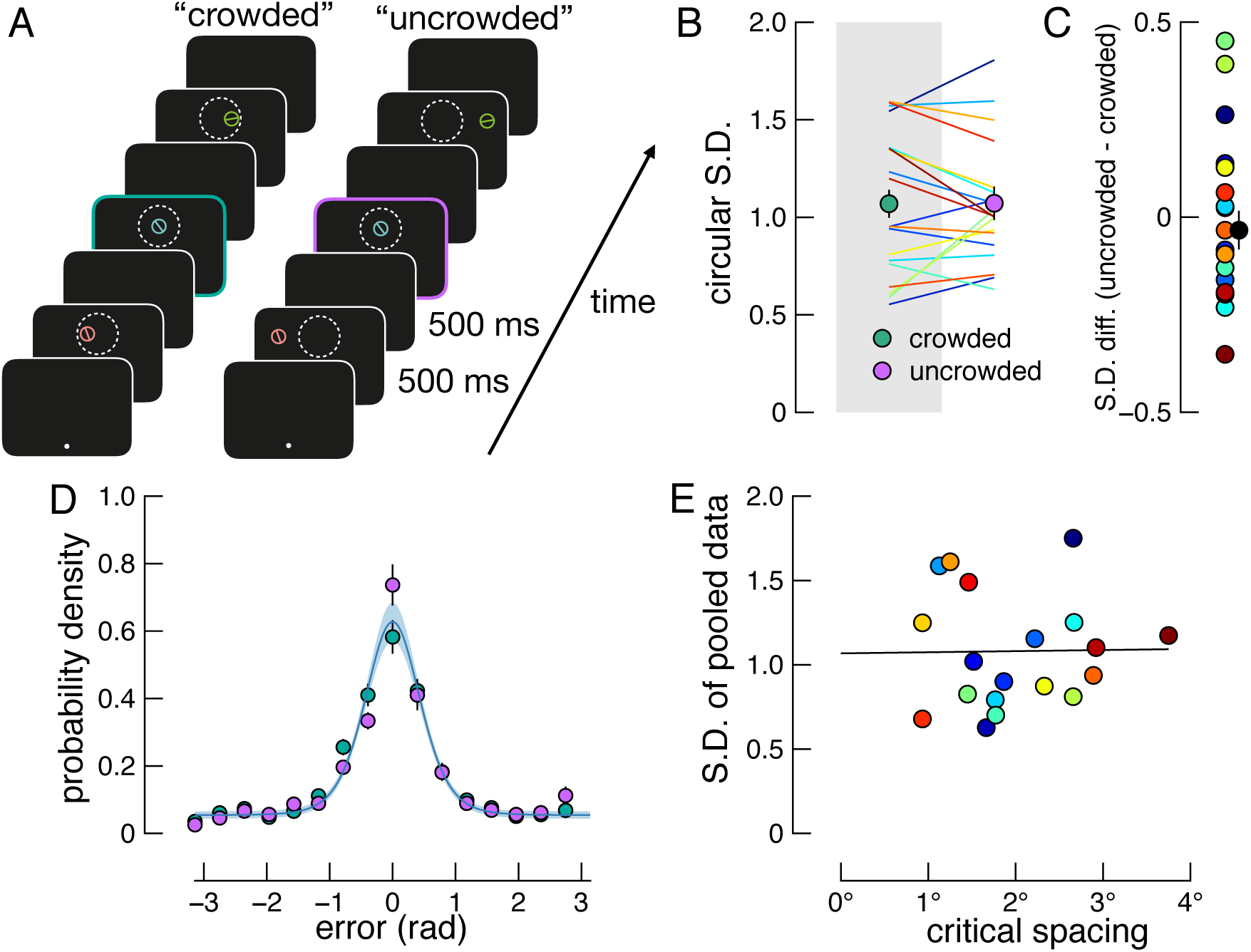
Experiment 2 design and results. A) Example trials of the crowded and uncrowded conditions. Observers fixated a white spot and viewed a sequence of randomly orientated memoranda that appeared within (crowded condition) or beyond (uncrowded condition) the critical spacing of their upper visual field, as indicated by the white dotted circle (shown for illustration only). After the final delay period, a probe appeared at one of the memorandum locations, and observers reported the target orientation at this location using a method of adjustment (see Methods). B) Report variability for each condition. Data are shown as in Fig. 2A. Coloured lines indicating each observer’s performance match colours in Figure 3C. C) Differences in report variability across conditions. Data are shown as in Fig. 2B. D) Error distributions and model fit. Green and purple points show crowded and uncrowded conditions, respectively. Data are shown as described in Fig. 2C. The model assuming memory performance is independent of cortical spacing (blue line) was again a better fit to the data than the model assuming an influence of cortical spacing, which has been omitted to increase visibility. E) Relationship between critical spacing and memory performance. We found no correlation between report variability pooled across conditions and critical spacing. Solid line indicates regression line of best fit.

#### Participants

21 participants took part in Experiment 2 (mean age 30.14 ± 8.69; 8 male, 13 female), one had also participated in Experiment 1, and all other details were as per the previous experiment. Two participants did not complete the experiment due to problems tracking their eyes, and their data were excluded from analyses, leaving a final sample size of 19.

#### Experimental Setup

All details were as per Experiment 1.

#### Stimuli

Stimuli were bars (0.85° × 0.04°) centred in a circle with a diameter matching the bar length, and a width of 0.04° (Fig. 3A and 4A). Three of such stimuli were displayed in each trial of both the crowding experiment and the memory experiment, and were uniquely coloured. We chose three colours equally spaced in CIE L*a*b* colour space, approximating red (L* = 74, a* = 34.6, b* = 20), green (L* = 74, a* = −28.3, b* = 28.3), and blue (L* = 74, a* = −28.3, b* = −28.3) hues. Colours were randomly assigned to the three stimuli on each trial. A white fixation spot was displayed in the centre of the screen throughout stimulus presentation and the memory delay period. All stimuli were presented on a black background.

In the crowding task, three oriented stimuli were presented simultaneously on each trial (Fig. 3A). The target orientation was random, while the distractors’ orientations were selected randomly from a uniform distribution that excluded orientations within 22.5° of the target orientation. Stimuli were centred 8.5° above fixation, and arranged tangentially relative to fixation. The centre stimulus was the target, and the others were distractors. As described below, the target-distractor distance was controlled via a staircase. Response stimuli were target and distractors in a neutral hue (grey), appearing in random positions but equally spaced on the border of an imaginary circle (radius = 1.7°) around the screen centre (Fig. 3A). When response stimuli were on screen, observers could move a standard mouse arrow that appeared in the screen centre. In the memory experiment, memoranda were of the same dimensions as the target and distractors in the crowding experiment, were each randomly assigned the colours described above, but were presented sequentially in random order. Stimulus orientations in the memory experiment were randomised with no restrictions.

#### Procedure

A typical trial sequence of the crowding task is shown in Fig. 3A. Each trial began following fixation compliance as per Experiment 1. Target and distractors appeared for 500ms. Following a 500ms delay, response stimuli and the response arrow appeared centered at fixation, and observers moved the arrow with the mouse and clicked on which stimulus they thought matched the target orientation. Observers were instructed that the target was always the central item on every trial, and that one response item matched its orientation exactly. No other instruction was explicitly given regarding the distractor response items, but if a participant asked about them, the experimenter told them that one item matched the target, and the other two response items matched the distractors. The next trial immediately followed each mouse click that fell within the border of a stimulus, and that stimulus was taken as their response.

The distance between the target and each distractor was controlled on each trial via an adaptive procedure, QUEST (Watson & Pelli, 1983), set to find the target-distractor spacing at which performance reached 67% accuracy (the midpoint of the psychometric function for a 3AFC task). We ran two randomly interleaved staircases of 36 trials each. For each QUEST procedure, we set the initial midpoint of the psychometric function (*μ*, see below) to two different levels to probe the asymptotes of the fitted function. These values, based on pilot observations, were set to 3.4° and 1.7°. These different QUEST parameters have the added advantage that the participant initially experiences relatively difficult and easy trials early on during testing. Furthermore, we allowed the target-distractor distance to vary only in steps of 0.21° during this threshold task. The procedure took approximately 7 minutes. Note that, while there was inevitably a working memory component to the crowding task, only the central element needed to be held in memory, therefore performance in this task indexes crowding occurring in sensory processing, due to the simultaneously presented flankers, rather than in memory.

The memory experiment was conducted in the same session as the crowding task, and is shown in Fig. 4A. Fixation compliance was performed, as above, and then each memory item was shown in random order, with a duration, inter-stimulus interval, and delay period of 500ms. Memoranda were shown in one of two spatial configurations, either spaced to fall within or beyond the critical spacing of crowding, as measured during the preceding task. After the delay period, a circle (diameter = 0.85°, width = 0.04°) matching the colour and location of one memory item was displayed, indicating to the observer to report that item’s orientation using the mouse. After the first mouse movement was registered, a response bar appeared within the circle so that the entire response stimulus matched the target dimensions. Observers then reported the target orientation as per Experiment 1 and the next trial began. Fixation errors and breaks were dealt with as described for Experiment 1. The crowding task and memory experiment took between 1 – 1.5 hours per observer. The number of trials per stimulus combination was as described in Experiment 1.

#### Experimental Design and Statistical Analyses

We pooled data across staircases in the crowded task and used the least-squares method to fit the Weibull function specified by Watson and Pelli (1983 see Fig. 3B). We modified the function to have three free parameters: μ, σ, and g, corresponding to the midpoint of the psychometric function, the slope, and the lapse rate, respectively. We took an observer’s critical spacing to be *μ*, which was bound between 0.85° and 8.5°, the lower of which ensured incomplete overlap of stimulus positions in the memory experiment for participants with very small crowding zones. Note that the lower bound was reached by only 2 out of 19 participants, while none reached the upper bound (Fig. 3C), and so this restriction is unlikely to have affected the results. The slope, *σ,* was bound between 0 and infinity, and lapse rate *g* was bound between 0 and 0.05 as recommended by Watson and Pelli (see also, Wichmann & Hill, 2001).

All comparisons in the memory experiment were within-subjects. We performed the same analyses of report variability and model fitting as per Experiment 1, but now with the conditions “crowded” and “uncrowded” to indicate trials in which memoranda were presented within or beyond the critical spacing of crowding, respectively. Importantly, these analyses were restricted to only memory items presented at the same screen position in both conditions so performance was matched in all aspects except for the spatial arrangement of memoranda. We further tested for a relationship between cortical spacing and short term memory with correlational analyses. We performed both a Bayesian Pearson Correlation and linear regression using JASP to test if memory performance, regardless of crowding level, could be predicted by critical spacing. We again restricted data to only trials in which the memory item was presented directly above fixation. For the Bayesian correlation, we used the default stretched beta prior width of 1, but results of this analysis were robust to various prior widths.

## Results

### Experiment 1

Perceptual resolution in peripheral vision is constrained by the distance between objects in primary visual cortex. As visual eccentricity increases, fewer visual neurons are available to process a constantly sized input, and this relationship is approximately logarithmic (Duncan & Boynton, 2003; Pelli, 2008). This log-scaling of visual cortex causes greater perceptual interference when multiple items are presented along a radial axis from the fovea compared with a tangential axis (Pelli & Tillman, 2008; Toet & Levi, 1992). In Experiment 1, we tested whether working memory is similarly influenced by the cortical spacing between memoranda (Fig. 1A). Observers were required to remember three sequentially presented randomly oriented bars arranged either radially or tangentially relative to the point of fixation (Fig. 1B). At the end of each sequence, observers’ memory of orientation was tested for a single item indicated by a location and colour probe (Fig. 1C), and responses were made by manually adjusting a response bar to match the cued item. To control for non-memory related differences across conditions, such as visual acuity, we analysed memory performance only for targets positioned at 10° to the right of fixation in each condition. These stimuli were matched in all regards except their spatial context.

Figure 2 summarises observers’ report errors for memoranda presented within a radial or tangential spatial configuration. As shown in Figure 2A, the circular standard deviation did not consistently differ between configurations. Indeed, a Bayesian paired-samples t-test found weak-to-moderate evidence in favour of there being no difference between conditions (B_01_ = 2.97; t(9) = 0.45, p = 0.66). These data provide evidence against the hypothesis that short term memory is worse when memoranda are more closely spaced in visual cortex.

**Figure 2.**
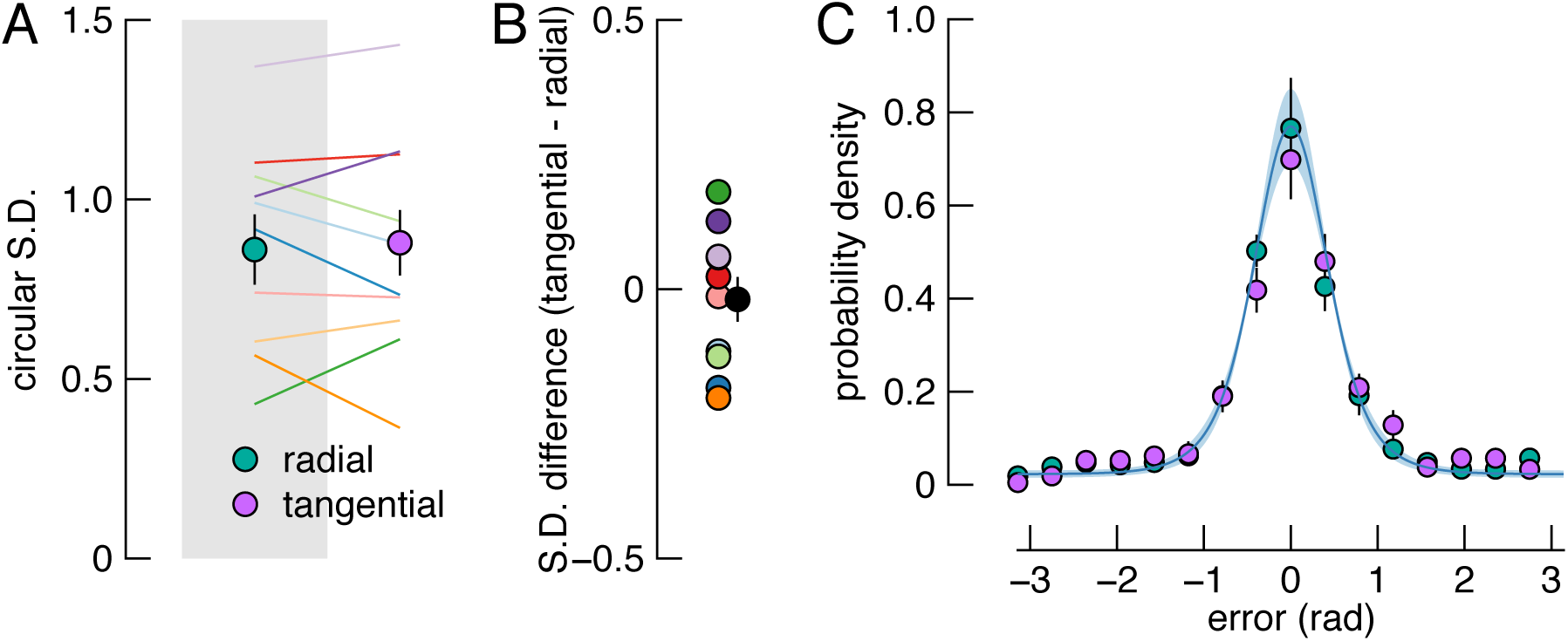
Results of Experiment 1. A) Report variability for each condition. Filled circles show the mean circular standard deviation of reports for radial (green) and tangential (purple) configurations. Coloured lines show individual participants’ data. Error bars indicate ±1 SE. B) Differences in report variability across conditions. The black datum shows the mean difference and the coloured data show individual difference scores, with colours corresponding to lines in (A). C) Error distributions and model fit. Frequency of errors for the radial and tangential conditions are expressed as probability densities, with colours as in (A). Data are shown for 16 equally spaced bins, in the range [−π π]. The solid blue line shows predictions of the best-fitting model, in which we assume memory is independent of the configuration of stimuli (shaded area indicates ±1 SE).

Figure 2C shows the distribution of errors in each condition. The solid line shows the fit of a model in which we assumed memory performance factors are independent of the arrangement of stimuli. This model was a better fit to the data than the model in which cortical spacing could influence memory performance (summed ΔAIC = 29.5; 8 out of 10 participants; Bayesian paired-samples t-test: B_10_ = 3.60; t(9) = 2.83, p = 0.02; maximum likelihood (ML) parameter values, mean (SE): precision = 5.21 (0.25); swaps = 0.10 (0.01); guesses = 0.14 (0.02)). This analysis further supports a dissociation between intra-cortical spacing and memory performance.

Finally, we ruled out the possibility that, although memory for the central item was unaffected, cortical spacing may have influenced the flanking memoranda which were excluded from the preceding analyses. We therefore repeated the above analyses, but included only trials in which the probed item was not in the central position. We first collated data across the remaining probe locations for each condition. We again found that there was no difference in circular standard deviation between radial and tangential conditions (B_01_ = 2.89; t(9) = 0.52, p = 0.62). The model in which we assume working memory is independent of cortical spacing was also the superior model (summed ΔAIC = 34.4; 9 out of 10 participants).

### Experiment 2

In Experiment 1 we manipulated intra-cortical spacing of memoranda by presenting items along a radial or tangential visual axis relative to fixation. We found positive evidence that performance was the same across conditions (Fig. 2A). These results suggest that visual short term memory does not have the properties of visual crowding that characterize retinotopic sensory areas that encode features. It is possible, however, that the stimulus arrangements we selected were not appropriately scaled to produce overlapping cortical mnemonic representations. To address this possibility, we conducted a second experiment in which we used a psychophysical approach to tailor intra-cortical spacing of memoranda individually for each participant.

We tested whether the cortical spacing of memoranda affects short term memory by sequentially presenting items either within or beyond the critical spacing of crowding. Critical spacing was found for each participant in a perceptual crowding task in which we used an adaptive staircase to find the target-distractor distance at which they could identify a target orientation at threshold level (Fig. 3A). Results from two example participants who performed differently at this task are shown in Figure 3B. Figure 3C shows the critical spacing estimates for all observers and the median for the group. Critical spacing estimates span an almost-fourfold range, and such between-subjects variability has been reported previously (Greenwood, Szinte, Sayim, & Cavanagh, 2017; Petrov & Meleshkevich, 2011). To control for between-subjects crowding variability in the memory experiment, and therefore control for cortical spacing variability across participants, we adjusted the spatial range of memoranda in the memory experiment to be either 0.75 times (“crowded”) or 1.5 times (“uncrowded”) an observer’s critical spacing.

Figure 4D shows the distribution of report errors averaged across observers, with green and purple data showing crowded and uncrowded conditions, respectively. We tested whether memory performance across conditions is better described by a model in which cortical spacing influences performance, or a model in which working memory is independent of cortical spacing of memoranda. The blue line in Figure 4D shows the model that is independent of cortical spacing, which was a better fit than the alternate model (summed ΔAIC = 52.46; 16 out of 19 participants; Bayesian paired-samples t-test: B_10_ = 150.2; t(18) = 2.83, p < 0.001; ML parameter values, mean (SE): precision = 5.62 (0.16); swaps = 0.04 (0.002); guesses = 0.34 (0.01)). Note that, although there is a higher probability density of uncrowded trials than crowded trials in the central bin (Fig. 4D; BF_10_ = 6.61), 16 bins were arbitrarily chosen for display purposes, and there would have been evidence against such a difference between conditions had we selected, for example, 15 bins (BF_10_ = 0.43). The analysis of variability and model fitting above are based on raw (unbinned) data so are not influenced by arbitrary designation of bin size.

Figure 4E shows the results of the correlational analysis in which we investigated whether there was a relationship between observers’ critical spacing and memory performance. A Bayesian Correlation Pairs test found moderate evidence that there is no relationship (r = 0.015, BF_01_ = 3.52). Similarly, a linear regression that uses critical spacing to predict report error found a slope of only 0.007 (t = 0.062, p = 0.951), indicating that there is no relationship between critical spacing and working memory performance.

As with Experiment 1, we again ruled out the possibility that cortical spacing may have influenced the flanking memoranda which were excluded from the preceding analyses. We repeated the above analyses including only trials in which the probed item was not in the central position, collapsing data across the remaining probe locations for each condition. In support of the results above, we found that there was no difference in circular standard deviation between crowded and uncrowded conditions (B_01_ = 4.08; t(18) = 0.27, p = 0.79). Finally, the model in which we assume working memory is independent of cortical spacing was superior (summed ΔAIC = 51.86; 16 out of 19 participants).

## Discussion

We investigated whether the cortical spacing between sequentially presented memoranda affects observers’ ability to hold those items in memory. In Experiment 1, we manipulated intra-cortical spacing by arranging memoranda either radially or tangentially relative to the fovea (Fig. 1). In Experiment 2, we tailored the intra-cortical spacing of memoranda to each observer by first quantifying their critical spacing of crowding (Fig. 3), and we then presented memory items within or beyond this region (Fig. 4). Across both experiments, we found positive evidence that working memory performance is independent of the cortical distance between memoranda. Although the strength of evidence in each experiment was only moderate, the combined evidence across experiments is assessed by the product of the individual Bayes Factors, i.e. 12.5, which is substantial.

Our study provides clear evidence of a dissociation between perceptual coding and memory coding within a very short period after stimulus offset. Cortical distance in retinotopically organised visual cortex can account for a wide variety of perceptual phenomena, such as visual acuity (Duncan & Boynton, 2003), shape perception (Michel, Chen, Geisler, & Seidemann, 2013), subjective experience of size (Schwarzkopf et al., 2011), and visual crowding (Pelli, 2008). In the present study, however, we have shown that memory representations of non-spatial features are independent of their V1 sensory representations. We know from our data that the emergence of dissociated representations occurs within the timeframe of the target duration and inter-stimulus interval (1 s). This time-course thus places an upper bound on the transfer of retinotopic sensory representations to other neural systems involved in working memory.

This result sheds light on previous psychophysical studies that have found errors in working memory due to spatially proximal memoranda. Pertzov et al (2014) and Ahmad et al (2017) found that memory for non-spatial features was worse when memoranda were presented sequentially at overlapping or similar screen locations than when memoranda were presented at spatially separate screen locations. However, the timing used in these experiments would have likely produced perceptual interference sometimes referred to as “temporal crowding” (Yeshurun et al., 2015). Such perceptual interference would degrade the encoding of memoranda due to their persistent overlapping cortical representations. Indeed, the nature of errors in these previous studies of working memory are consistent with those in visual crowding paradigms with minimal working memory demands (Ester, Klee, & Awh, 2014; Harrison & Bex, 2015; 2017). The combination of target duration and interstimulus interval used by Pertzov et al. (500 ms) thus sets a lower bound on the time required to transform a sensory signal into a memory representation.

Our results raise several important challenges for the hypothesis that working memory representations are maintained via the same sensory neurons that encoded the features of memoranda (Serences, 2016). Previous studies in which a remembered feature is decoded from activity within V1 typically analyse activity within voxels corresponding to the spatial location of the memory item (e.g. Harrison & Tong, 2009; Serences et al., 2009). Because our data reveal that sensory representations are independent of memory representations, these decoding analyses must either be decoding non-sensory neurons that are nonetheless tuned to the memoranda feature dimension, which we think is unlikely, or reflect an influence from other areas. Other brain regions implicated in memory maintenance include prefrontal cortex and posterior parietal cortex (Bettencourt & Xu, 2016; Christophel, Hebart, & Haynes, 2012; Courtney, Petit, Maisog, Ungerleider, & Haxby, 1998; Todd & Marois, 2004). In prefrontal cortex in particular, neurons display activity during memory delays that encodes stimulus locations and features (Goldman-Rakic, 1995; Mendoza-Halliday & Martinez-Trujillo, 2017; Murray et al., 2017; Wimmer, Nykamp, Constantinidis, & Compte, 2014) (but see Lara & Wallis, 2014). These areas are part of a distributed network involved in working memory, and the role of V1 in this network remains to be fully understood (Christophel, Klink, Spitzer, Roelfsema, & Haynes, 2017; D’Esposito, 2007; D’Esposito & Postle, 2015).

Another alternative is that working memory is maintained via the recruitment of sensory neurons well beyond the initial sensory representation (Ester, Serences, & Awh, 2009). According to this *neural outsourcing* proposal, the memory representation of a stimulus might be shifted to neurons that normally encode sensory stimulation in some other part of the visual field. However, it is yet to be clarified how visual features with overlapping sensory representations are allocated to other sensory regions in a way that prevents memory interference, nor how a mapping is maintained between outsourced representations and their original locations in the visual field.

Bays (2014) recently proposed a neural resource model of working memory, based on population coding, that can account for changes in memory precision as a function of the number of memoranda. A key feature of this model is that a fixed amount of neural activity (i.e. spiking) must be shared amongst all memory items. Increasing set size, therefore, decreases the neural resource available for each item, resulting in a loss of memory precision. The property of maintaining a fixed level of population activity is termed normalisation: it has been described as a canonical neural computation, implemented in many different neural systems using varied mechanisms (Carandini & Heeger, 2012). To accurately reproduce observed effects of set size, the normalisation in the model must operate globally, i.e. not limited to particular regions of the visual field or particular stimulus feature values (Bays, 2015). The present results are in agreement with this, in that they confirm there is no cost of spatial proximity of memoranda as might be expected from a purely local form of normalisation.

Neurophysiological evidence consistent with global normalisation has been found in prefrontal and posterior parietal cortices, areas which have been implicated as playing an important role in working memory maintenance (for a review, see Bays, 2015). Although inspired by properties of visual neurons, the neural resource model is agnostic as to the neural locus of working memory representations, as population coding is a common mechanism of representation observed throughout the brain (Pouget, Dayan, & Zemel, 2000), including prefrontal cortex (Murray et al., 2017; Wimmer et al., 2014). Nonetheless, one possible interpretation, consistent with the present findings, is that, in the case of visual working memory representations, normalisation occurs within networks in which neurons are not strictly topographically organised.

Although neural models of short term memory can account for a broad range of human performance, we are not aware of any model that can account for our result. In a recent study, Schneegans and Bays (2017) presented strong evidence in favour of a model in which non-spatial features are combined with spatial location via a conjunctive population code. This extension of the neural resource model correctly predicted their empirical observation that, when memoranda are presented simultaneously, observers were more likely to confuse items in working memory (“swap” errors) when the cued memory item was close to distractors than when distractors were relatively distant from the cued item.

This model is also consistent with the results of Tamber-Rosenau et al (2015), who found that the frequency of swap errors for simultaneously presented memoranda depends on the degree of perceptual crowding. Because visual crowding increases positional uncertainty (Harrison & Bex, 2017), a conjunctive code that binds spatial location with orientation will produce more swap errors under strongly-crowded conditions than weakly-crowded conditions, as was observed by Tamber-Rosenau et al. Schneegans and Bays’ (2017) model therefore suggests an important role of location in binding non-spatial features when items are presented simultaneously, but leaves open the question of how to account for the present findings with sequentially presented memoranda. It is possible that non-spatial features can be bound according to a conjunctive code that links features with their temporal order, but neurophysiological evidence for such a model is scarce. Accounting for the lack of spatial interactions between sequentially presented memoranda represents a challenge for future modelling efforts.

## Acknowledgments

This work was supported by the Wellcome Trust, and a National Health and Medical Research Council of Australia CJ Martin Fellowship to W.J.H. (APP1091257).

## Author Contributions

W.J.H. and P.M.B. designed research; W.J.H. performed research; W.J.H analysed data; W.J.H. and P.M.B. wrote the paper.

## References

Ahmad, J., Swan, G., Bowman, H., Wyble, B., Nobre, A. C., Shapiro, K. L., & McNab, F. (2017). Competitive interactions affect working memory performance for both simultaneous and sequential stimulus presentation. Scientific Reports, 7(1), 4785. http://doi.org/10.1038/s41598-017-05011–x

Anderson, E. J., Dakin, S. C., Schwarzkopf, D. S., Rees, G., & Greenwood, J. A. (2012). The neural correlates of crowding-induced changes in appearance. Current Biology, 22(13). http://doi.org/10.1016/j.cub.2012.04.063

Bays, P. M. (2014). Noise in Neural Populations Accounts for Errors in Working Memory. The Journal of Neuroscience, 34(10), 3632–3645. http://doi.org/10.1523/JNEUROSCI.3204-13.2014

Bays, P. M. (2015). Spikes not slots: noise in neural populations limits working memory. Trends in Cognitive Sciences, 19(8), 431–438. http://doi.org/10.1016/j.tics.2015.06.004

Bays, P. M., Catalao, R. F. G., & Husain, M. (2009). The precision of visual working memory is set by allocation of a shared resource. Journal of Vision, 9(10), 7. 1–11. http://doi.org/10.1167/9.10.7

Bettencourt, K. C., & Xu, Y. (2016). Decoding the content of visual short-term memory under distraction in occipital and parietal areas. Nature Neuroscience, 19(1), 150–. http://doi.org/10.1038/nn.4174

Brainard, D. H. (1997). The Psychophysics Toolbox. Spatial Vision, 10(4), 433–436.

Carandini, M., & Heeger, D. J. (2012). Normalization as a canonical neural computation. Nature Reviews Neuroscience, 13(1), 51–62. http://doi.org/10.1038/nrn3136

Chen, J., He, Y., Zhu, Z., Zhou, T., Peng, Y., Zhang, X., & Fang, F. (2014). Attention-dependent early cortical suppression contributes to crowding. Journal of Neuroscience, 34(32), 10465–10474. http://doi.org/10.1523/JNEUROSCI.1140-14.2014

Christophel, T. B., Hebart, M. N., & Haynes, J.-D. (2012). Decoding the contents of visual short-term memory from human visual and parietal cortex. Journal of Neuroscience, 32(38), 12983–12989. http://doi.org/10.1523/JNEUROSCI.0184-12.2012

Christophel, T. B., Klink, P. C., Spitzer, B., Roelfsema, P. R., & Haynes, J.-D. (2017). The Distributed Nature of Working Memory. Trends in Cognitive Sciences, 21(2), 111–124. http://doi.org/10.1016/j.tics.2016.12.007

Cornelissen, F. W., Peters, E. M., & Palmer, J. (2002). The Eyelink Toolbox: eye tracking with MATLAB and the Psychophysics Toolbox. Behavior Research Methods, Instruments & Computers, 34(4), 613–617.

Courtney, S. M., Petit, L., Maisog, J. M., Ungerleider, L. G., & Haxby, J. V. (1998). An area specialized for spatial working memory in human frontal cortex. Science, 279(5355), 1347–1351. http://doi.org/10.1126/science.279.5355.1347

D'Esposito, M. (2007). From cognitive to neural models of working memory. Philosophical Transactions of the Royal Society of London. Series B, Biological Sciences, 362(1481), 761–772. http://doi.org/10.1098/rstb.2007.2086

D'Esposito, M., & Postle, B. R. (2015). The cognitive neuroscience of working memory. Annual Review of Psychology, 66, 115–142. http://doi.org/10.1146/annurev-psych-010814-015031

Duncan, R. O., & Boynton, G. M. (2003). Cortical magnification within human primary visual cortex correlates with acuity thresholds. Neuron, 38(4), 659–671.

Emrich, S. M., Riggall, A. C., LaRocque, J. J., & Postle, B. R. (2013). Distributed Patterns of Activity in Sensory Cortex Reflect the Precision of Multiple Items Maintained in Visual Short-Term Memory. Journal of Neuroscience, 33(15), 6516–6523. http://doi.org/10.1523/JNEUROSCI.5732-12.2013

Ester, E. F., Klee, D., & Awh, E. (2014). Visual crowding cannot be wholly explained by feature pooling. Journal of Experimental Psychology: Human Perception and Performance, 40(3), 1022–1033. http://doi.org/10.1037/a0035377

Ester, E. F., Serences, J. T., & Awh, E. (2009). Spatially global representations in human primary visual cortex during working memory maintenance. Journal of Neuroscience, 29(48), 15258–15265. http://doi.org/10.1523/JNEUROSCI.4388-09.2009

Freeman, J., & Simoncelli, E. P. (2011). Metamers of the ventral stream. Nature Neuroscience, 14(9), 1195–1201. http://doi.org/10.1038/nn.2889

Goldman-Rakic, P. S. (1995). Cellular basis of working memory. Neuron, 14(3), 477–485.

Gorgoraptis, N., Catalao, R. F. G., Bays, P. M., & Husain, M. (2011). Dynamic updating of working memory resources for visual objects. Journal of Neuroscience, 31(23), 8502–8511. http://doi.org/10.1523/JNEUROSCI.0208-11.2011

Greenwood, J. A., Szinte, M., Sayim, B., & Cavanagh, P. (2017). Variations in crowding, saccadic precision, and spatial localization reveal the shared topology of spatial vision. Proceedings of the National Academy of Sciences of the United States of America, 23, 201615504. http://doi.org/10.1016/0042-6989(72)90090-9

Harrison, S. A., & Tong, F. (2009). Decoding reveals the contents of visual working memory in early visual areas. Nature, 458(7238), 632–635. http://doi.org/10.1038/nature07832

Harrison, W. J., & Bex, P. J. (2015). A Unifying Model of Orientation Crowding in Peripheral Vision. Current Biology, 25(24), 3213–3219. http://doi.org/10.1016/j.cub.2015.10.052

Harrison, W. J., & Bex, P. J. (2017). Visual crowding is a combination of an increase of positional uncertainty, source confusion, and featural averaging. Scientific Reports, 7, 45551. http://doi.org/10.1038/srep45551

Kwon, M., Bao, P., Millin, R., & Tjan, B. S. (2014). Radial-tangential anisotropy of crowding in the early visual areas. Journal of Neurophysiology, 112(10), 2413–2422. http://doi.org/10.1152/jn.00476.2014

Lara, A. H., & Wallis, J. D. (2014). Executive control processes underlying multi-item working memory. Nature Neuroscience, 17(6), 876–883. http://doi.org/10.1038/nn.3702

McGraw, P. V., & Whitaker, D. (1999). Perceptual distortions in the neural representation of visual space. Experimental Brain Research, 125(2), 122–128. http://doi.org/10.1007/s002210050667

Mendoza-Halliday, D., & Martinez-Trujillo, J. C. (2017). Neuronal population coding of perceived and memorized visual features in the lateral prefrontal cortex. Nature Communications, 8, 15471. http://doi.org/10.1038/ncomms15471

Michel, M. M., Chen, Y., Geisler, W. S., & Seidemann, E. (2013). An illusion predicted by V1 population activity implicates cortical topography in shape perception. Nature Neuroscience, 16(10), 1477–1483. http://doi.org/10.1038/nn.3517

Murray, J. D., Bernacchia, A., Roy, N. A., Constantinidis, C., Romo, R., & Wang, X.-J. (2017). Stable population coding for working memory coexists with heterogeneous neural dynamics in prefrontal cortex. Proceedings of the National Academy of Sciences of the United States of America, 114(2), 394–399. http://doi.org/10.1073/pnas.1619449114

Pasternak, T., & Greenlee, M. W. (2005). Working memory in primate sensory systems. Nature Reviews Neuroscience, 6(2), 97–107. http://doi.org/10.1038/nrn1603

Pelli, D. G. (1997). The VideoToolbox software for visual psychophysics: Transforming numbers into movies. Spatial Vision, 10(4), 437–442.

Pelli, D. G. (2008). Crowding: a cortical constraint on object recognition. Current Opinion in Neurobiology, 18(4), 445–451. http://doi.org/10.1016Zj.conb.2008.09.008

Pelli, D. G., & Tillman, K. A. (2008). The uncrowded window of object recognition. Nature Neuroscience, 11(10), 1129–1135.

Pertzov, Y., & Husain, M. (2014). The privileged role of location in visual working memory. Attention, Perception & Psychophysics, 76(7), 1914–1924. http://doi.org/10.3758/s13414-013-0541-y

Petrov, Y., & Meleshkevich, O. (2011). Asymmetries and idiosyncratic hot spots in crowding. Vision Research, 51(10), 1117–1123. http://doi.org/10.1016Zj.visres.2011.03.001

Pouget, A., Dayan, P., & Zemel, R. (2000). Information processing with population codes. Nature Reviews Neuroscience, 1(2), 125–132. http://doi.org/10.1038/35039062

Schneegans, S., & Bays, P. M. (2017). Neural Architecture for Feature Binding in Visual Working Memory. Journal of Neuroscience, 37(14), 3913–3925. http://doi.org/10.1523/JNEUROSCI.3493-16.2017

Schwarzkopf, D. S., Song, C., & Rees, G. (2011). The surface area of human V1 predicts the subjective experience of object size. Nature Neuroscience, 14(1), 28–30. http://doi.org/10.1038/nn.2706

Serences, J. T. (2016). Neural mechanisms of information storage in visual short-term memory. Vision Research, 128, 53–67. http://doi.org/10.1016/j.visres.2016.09.010

Serences, J. T., Ester, E. F., Vogel, E. K., & Awh, E. (2009). Stimulus-specific delay activity in human primary visual cortex. Psychological Science, 20(2), 207–214. http://doi.org/10.1111/j.1467-9280.2009.02276.x

Sprague, T. C., Ester, E. F., & Serences, J. T. (2014). Reconstructions of information in visual spatial working memory degrade with memory load. Current Biology, 24(18), 2174–2180. http://doi.org/10.1016/j.cub.2014.07.066

Sreenivasan, K. K., Curtis, C. E., & D'Esposito, M. (2014). Revisiting the role of persistent neural activity during working memory. Trends in Cognitive Sciences, 18(2), 82–89. http://doi.org/10.1016/j.tics.2013.12.001

Tamber-Rosenau, B. J., Fintzi, A. R., & Marois, R. (2015). Crowding in Visual Working Memory Reveals Its Spatial Resolution and the Nature of Its Representations. Psychological Science. http://doi.org/10.1177/0956797615592394

Todd, J. J., & Marois, R. (2004). Capacity limit of visual short-term memory in human posterior parietal cortex. Nature, 428(6984), 751–754. http://doi.org/10.1038/nature02466

van den Berg, R., Roerdink, J. B. T. M., & Cornelissen, F. W. (2010). A neurophysiologically plausible population code model for feature integration explains visual crowding. PLoS Computational Biology, 6(1), e1000646. http://doi.org/10.1371/journal.pcbi.1000646

Watson, A. B., & Pelli, D. G. (1983). QUEST: a Bayesian adaptive psychometric method. Perception and Psychophysics, 33(2), 113–120.

White, J. M., Levi, D. M., & Aitsebaomo, A. P. (1992). Spatial localization without visual references. Vision Research. http://doi.org/10.1016/0042-6989(92)90243-C

Wichmann, F. A., & Hill, N. J. (2001). The psychometric function: I. Fitting, sampling, and goodness of fit. Attention, Perception & Psychophysics, 63(8), 1293–1313.

Wimmer, K., Nykamp, D. Q., Constantinidis, C., & Compte, A. (2014). Bump attractor dynamics in prefrontal cortex explains behavioral precision in spatial working memory. Nature Neuroscience, 17(3), 431–439. http://doi.org/10.1038/nn.3645

Xu, Y. (2017). Reevaluating the Sensory Account of Visual Working Memory Storage. Trends in Cognitive Sciences. http://doi.org/10.1016/j.tics.2017.06.013

Yeshurun, Y., Rashal, E., & Tkacz-Domb, S. (2015). Temporal crowding and its interplay with spatial crowding. Journal of Vision, 15(3). http://doi.org/10.1167/15.3.11

Zhang, W., & Luck, S. J. (2008). Discrete fixed-resolution representations in visual working memory. Nature, 453(7192), 233–235. http://doi.org/10.1038/nature06860

